# Memory Th1 cells modulate heterologous diseases through innate function

**DOI:** 10.1101/2023.03.22.533799

**Authors:** Nikolas Rakebrandt, Nima Yassini, Anna Kolz, Michelle Schorer, Katharina Lambert, Celine Rauld, Zsolt Balazs, Michael Krauthammer, José M. Carballido, Anneli Peters, Nicole Joller

**Author notes:** These authors contributed equally to this work (Nikolas Rakebrandt, Nima Yassini).

## Abstract

Through immune memory, infections have a lasting effect on the host. While memory cells enable accelerated and enhanced responses upon re-challenge with the same pathogen, their impact on susceptibility to unrelated diseases is unclear. We identify a subset of memory T helper 1 (Th1) cells termed innate acting memory T (T_IA_) cells that originate from a viral infection and produce IFN-γ with innate kinetics upon heterologous challenge *in vivo*. Activation of memory T_IA_ cells is induced in response to IL-12 in combination with IL-18 or IL-33 but is TCR-independent. Rapid IFN-γ production by memory T_IA_ cells is protective in subsequent heterologous challenge with the bacterial pathogen *Legionella pneumophila*. In contrast, antigen-independent re-activation of CD4^+^ memory T_IA_ cells accelerates disease onset in an autoimmune model of multiple sclerosis. Our findings demonstrate that memory Th1 cells can acquire additional TCR-independent functionality to mount rapid, innate-like responses that modulate susceptibility to heterologous challenges.

## Introduction

Disease susceptibility can be highly variable between individuals as highlighted by the broad range of disease courses seen e.g. in the recent COVID-19 pandemic (Arunachalam et al., 2020; Mathew et al., 2020; Zhou et al., 2020). Genetic predisposition influences the propensity of the immune system to respond to challenges as well as the magnitude of that response. In addition, environmental factors as well as interactions with pathogens and the microbiome contribute to the variability observed in disease susceptibility (Orrù et al., 2013; Brodin et al., 2015; Farh et al., 2015; Carr et al., 2016; Cheung et al., 2018; Ellinghaus et al., 2020). Pathogen exposure triggers a transient effector response but also establishes a persisting pool of memory cells, that play an essential role in mediating long-term protection against secondary infections with the same pathogen (Lanzavecchia and Sallusto, 2000). However, their impact on heterologous challenges is less clear. Although cross-reactive memory cells can alter the disease course in some settings (Lang et al., 2002; Bardina et al., 2017), their impact is restricted to very few specific combinations. Besides TCR-dependent activation, several *in vitro* studies have shown that memory T cells may also be activated in response to certain cytokine combinations through so called bystander activation (Geginat et al., 2001; Guo et al., 2009; Sattler et al., 2009; Freeman et al., 2012). Cytokine-mediated activation and IFN-γ production by CD8^+^ T cells is potently induced by IL-12 + IL-18 and to a lesser degree by IL-12 + TNFɑ (Raué et al., 2004; Freeman et al., 2012). Similarly, CD4^+^ T helper cells can be activated in the absence of a TCR trigger in response to combinations of a STAT-activator and an IL-1 family cytokine member (Guo et al., 2009).

Bystander activation of memory CD8^+^ T cells, has been shown to influence disease severity in several disease settings including rheumatoid arthritis, hepatitis and COVID-19 (Groh et al., 2003; Kim et al., 2018; Bergamaschi et al., 2021). However, whether and how memory CD4^+^ T helper cells established during prior infections influence the magnitude and more importantly the nature of subsequent immune responses to heterologous challenges *in vivo* is still unclear. In this study, we investigated the response of virus-specific memory Th1 cells in heterologous challenges. To induce memory, we performed acute infections with Lymphocytic Choriomeningitis virus (LCMV) and then re-challenged the mice with the unrelated bacterial pathogen *L. pneumophila* (Lpn) after the viral infection had been cleared. Even though the two pathogens do not harbor shared antigens, virus-specific memory CD4^+^ T cells mounted an early IFN-γ response upon bacterial challenge, which was sufficient to reduce the bacterial burden. The response of these innate acting memory T cells (T_IA_ cells) was TCR-independent and could be induced by cytokine stimulation alone. Furthermore, T_IA_ cells displayed a superior migratory capability that was essential for the protective effect observed in the bacterial challenge. In an autoimmune setting, the rapid, antigen-independent activation and enhanced migratory capacity of CD4^+^ T_IA_ cells enabled them to infiltrate the CNS and contribute to an earlier disease onset in a model of multiple sclerosis. Our findings thus uncovered a new facet of memory CD4^+^ T cells *in vivo*, whereby they bear the potential to respond rapidly to heterologous challenges in a TCR-independent fashion that ultimately alters disease severity.

## Results

### Heterologous protection from infection

To address if and how memory T cells influence the outcome of unrelated challenges, we established a heterologous infection model using two antigenically distinct pathogens. We observed that prior LCMV infection conferred partial protection from a later bacterial challenge with Lpn as indicated by reduced bacterial titers 3 days post challenge when compared to a control group (Figures 1A and 1B). Using high dimensional CyTOF analysis to compare cell composition and function in the lungs of memory and control mice, we observed an increase in the proportion of neutrophils, as well as T and B cells in memory mice and a shift in IFN-γ production from NK towards T cells (Figure S1A). Interestingly, a prominent IFN-γ response was evident in the absence of antigen-specific restimulation in both CD4^+^ and CD8^+^ T cells from memory but not control mice, which was confirmed by classical flow cytometry (Figures 1C, 1D, and S1A-C). These results are in line with previous studies reporting early bystander activation of CD8^+^ T cells in a number of settings (Groh et al., 2003; Freeman et al., 2012; Kim et al., 2018). In addition, we also detected robust IFN-γ production in CD4^+^ T cells upon heterologous challenge *in vivo*, where previous studies were limited to *in vitro* settings without prior infection (Nogai et al., 2005; Lee et al., 2019).

**Figure 1.**
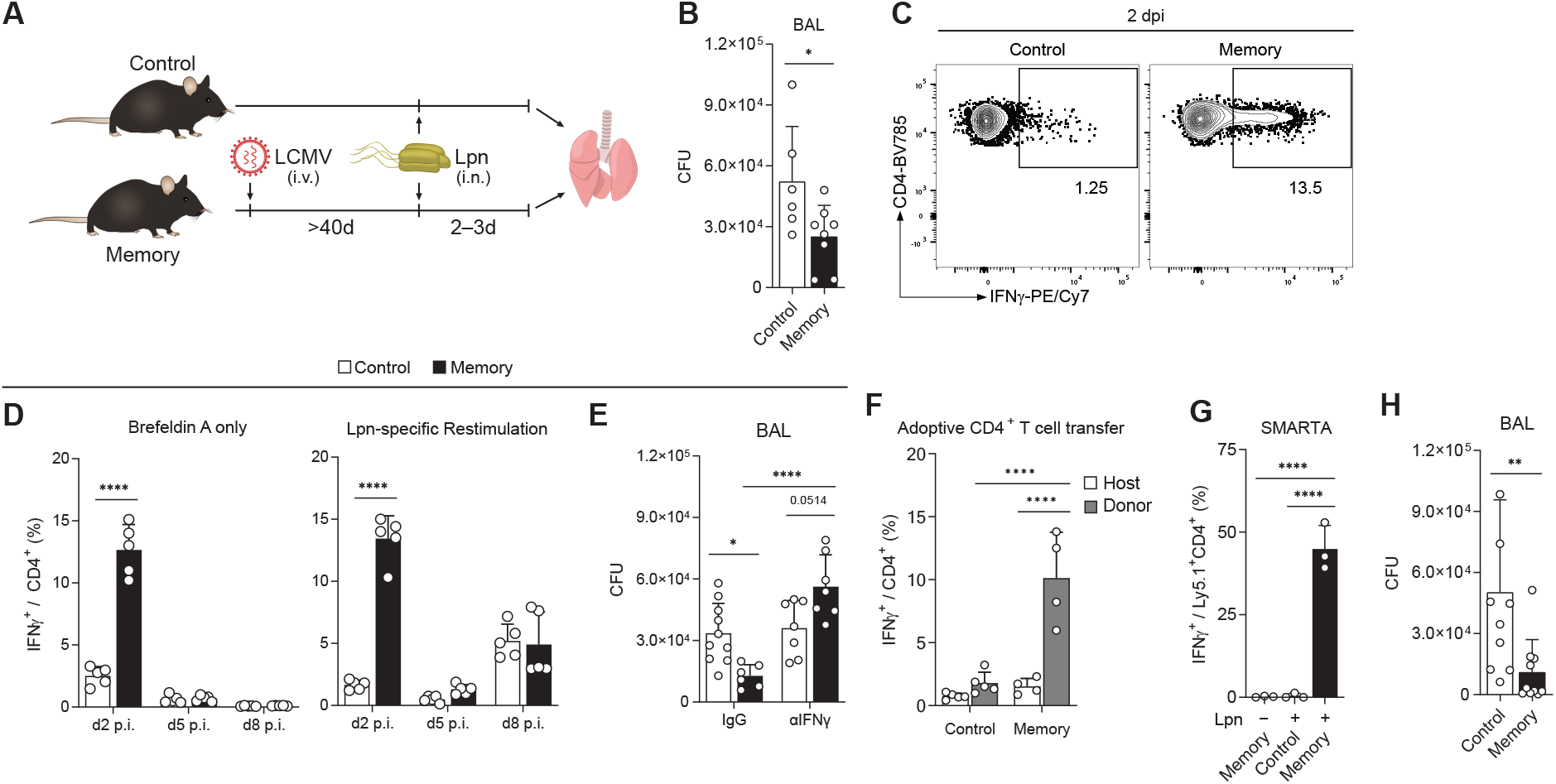
Early IFN-γ production in virus-experienced CD4^+^ T cells mediates protection against Lpn Control and LCMV-experienced memory mice were challenged with Lpn. **A**, Experimental layout of the heterologous infection model. **B**, Bacterial titers from bronchoalveolar lavage (BAL) fluid 3 days post infection (dpi.) (n=6–8). **C**, Representative FACS plot of the IFN-γ response of CD4^+^ T cells. **D**, Time course of IFN-γ response with or without Lpn restimulation (n=5). **E**, Bacterial titers 3 dpi. in mice treated intranasally on d0 and d1 with anti-IFN-γ neutralizing or IgG control antibody (n = 6–10). **F**, 5×10^5^ CD4^+^ T cells isolated from the spleen were transferred i.v. one day before Lpn infection into congenic hosts and the IFN-γ response was measured 60h post infection (n = 4–5). **G**, Mice received 5×10^4^ SMARTA cells i.v. one day prior to LCMV infection (memory) or 10^6^ naïve SMARTA cells one day before Lpn challenge (control). IFN-γ response was measured on day 2 post Lpn challenge (n = 3). **H**, Bacterial titers 3 dpi. of *Rag2*^−/−^g*c*^−/−^ mice that received 2×10^6^ naïve or memory CD4^+^ T cells 2 days prior to Lpn infection (n = 9–10). Mean ± s.d., unpaired t test (B), two-way (*Šídák*; D, E, F) or one-way (Tukey; G) ANOVA, and Mann-Whitney test (H).

Analysis of the T cell response over time revealed that, like CD8^+^ T cells, CD4^+^ T cells from memory mice readily produced IFN-γ as early as 2 days post infection, which declined again thereafter (Figures 1D and S1C). Importantly, this early IFN-γ production was observed in the absence of pathogen-specific restimulation with Lpn and did not require any T cell stimulating agents such as PMA/ionomycin or ɑCD3/ɑCD28 treatment (Figure 1D). Additionally, this innate-like response was distinct from the antigen-specific T cell response towards Lpn detected on day 8, which was comparable between control and memory mice (Figures 1D and S1C). Furthermore, antigen-independent cytokine production was restricted to IFN-γ as neither TNF-ɑ nor IL-17 could be detected at these early timepoints or in the absence of antigen-specific stimulation (Figures S1C and S1D). This suggested that the early IFN-γ peak was dependent on memory Th cells generated during the earlier virus infection, while the second wave of IFN-γ was likely elicited by *de novo* priming upon Lpn infection. To confirm that IFN-γ is protective in this disease setting as previously reported (Brown et al., 2016), we blocked IFN-γ upon Lpn challenge and found that this abolished the heterologous protection (Figure 1E), supporting the notion that the early antigen-independent IFN-γ secretion from memory T cells mediates the protective effect observed during the heterologous challenge.

To further investigate the mechanism of antigen-independent reactivation of CD4^+^ T cells, we first confirmed that the ability to rapidly produce IFN-γ upon heterologous challenge was not pathogen-specific and limited to LCMV-specific memory CD4^+^ T cells, as it also occurred after an initial vaccinia virus infection (Figure S1E). Rapid IFN-γ production upon heterologous challenge thus represents a common feature of memory Th1 cells. To determine whether this altered IFN-γ response was the result of a virus-experienced environment enabling the rapid cytokine production or of T cell-intrinsic features, we transferred memory (or control) CD4^+^ T cells into naïve hosts before Lpn challenge and analyzed the innate-like response. This adoptive transfer revealed that the ability for early IFN-*γ* production was T cell-intrinsic as transferred memory but not control or endogenous CD4^+^ T cells secreted IFN-*γ* upon Lpn challenge (Figures 1F and S1F). Furthermore, transfer of Smarta CD4^+^ T cells, which carry a TCR specific for the LCMV gp61 peptide and do not react to Lpn antigens (Figure S1G), confirmed that IFN-γ production was not a consequence of cross-reactivity, since memory but not control Smarta T cells produced high amounts of IFN-γ 2 days after Lpn challenge and returned to baseline by day 5 (Figures 1G and S1H). Importantly, although memory CD8^+^ T cells also rapidly produced IFN-γ upon heterologous challenge (Figure S1C), transfer of CD4^+^ T cells from control vs. memory mice into *Rag2*^−/−^g*c*^−/−^ mice lacking T, B, and NK cells was sufficient to replicate the heterologous protection observed (Figure 1H), highlighting their functional contribution. Viral infections therefore induce a CD4^+^ memory Th1 cell population that can mount a rapid antigen-independent IFN-γ response. These innate acting memory CD4^+^ T cells (T_IA_ cells) can mediate heterologous protection by antigen-independent, early IFN-γ production when faced with an unrelated pathogenic challenge.

### Innate acting memory CD4^+^ T cells

To better characterize these CD4^+^ T_IA_ cells, we revisited our initial CyTOF analysis to look for markers that may distinguish CD4^+^ T_IA_ cells producing IFN-*γ* upon heterologous challenge from other memory CD4^+^ T cells. Such distinction would enable us to identify CD4^+^ T_IA_ cells at steady state even before they start secreting IFN-γ. Indeed, compared to IFN-γ^−^ CD4^+^ T cells, CD4^+^ TIA cells showed higher expression of the germline-encoded receptor NKG2D (Figure S2A). NKG2D expression on CD4^+^ T cells has been associated with autoimmune disorders in mice and humans (Groh et al., 2003; Ruck et al., 2013; Yang et al., 2017) and has been linked to bystander activation on CD8^+^ T cells (Whiteside et al., 2018). Classical flow cytometry confirmed that the NKG2D^+^CD4^+^ T cell fraction was highly enriched for IFN-γ^+^ cells and we found the CD44^+^NKG2D^+^CD4^+^ T cell fraction to be strongly expanded in memory mice (Figures 2A and S2B), indicating that NKG2D can be used as a marker for CD4^+^ T_IA_ cells. We next compared the numbers of NKG2D^+^CD4^+^ T cells before and after Lpn challenge to determine whether CD4^+^ T_IA_ cells are already present in the lung before challenge or actively recruited to the site of infection. Memory, but not control mice showed a marked increase in NKG2D^+^CD4^+^ T cells in the lung accompanied by a decline in the spleen upon heterologous Lpn challenge (Figures S2C and S2D), suggesting that CD4 T_IA_ cells are recruited from the spleen to the lung upon challenge. Indeed, blockade of T cell migration from secondary lymphoid organs using Fingolimod (FTY720) inhibited the increase in NKG2D^+^CD4^+^ T cell numbers in the lung and significantly reduced the number of IFN-γ^+^ CD4^+^ T cells upon Lpn challenge (Figures S2E and S2F). The spleen thus appears to represent a reservoir for NKG2D^+^CD4^+^ T cells, which are recruited to peripheral sites upon heterologous challenge to produce IFN-γ.

**Figure 2.**
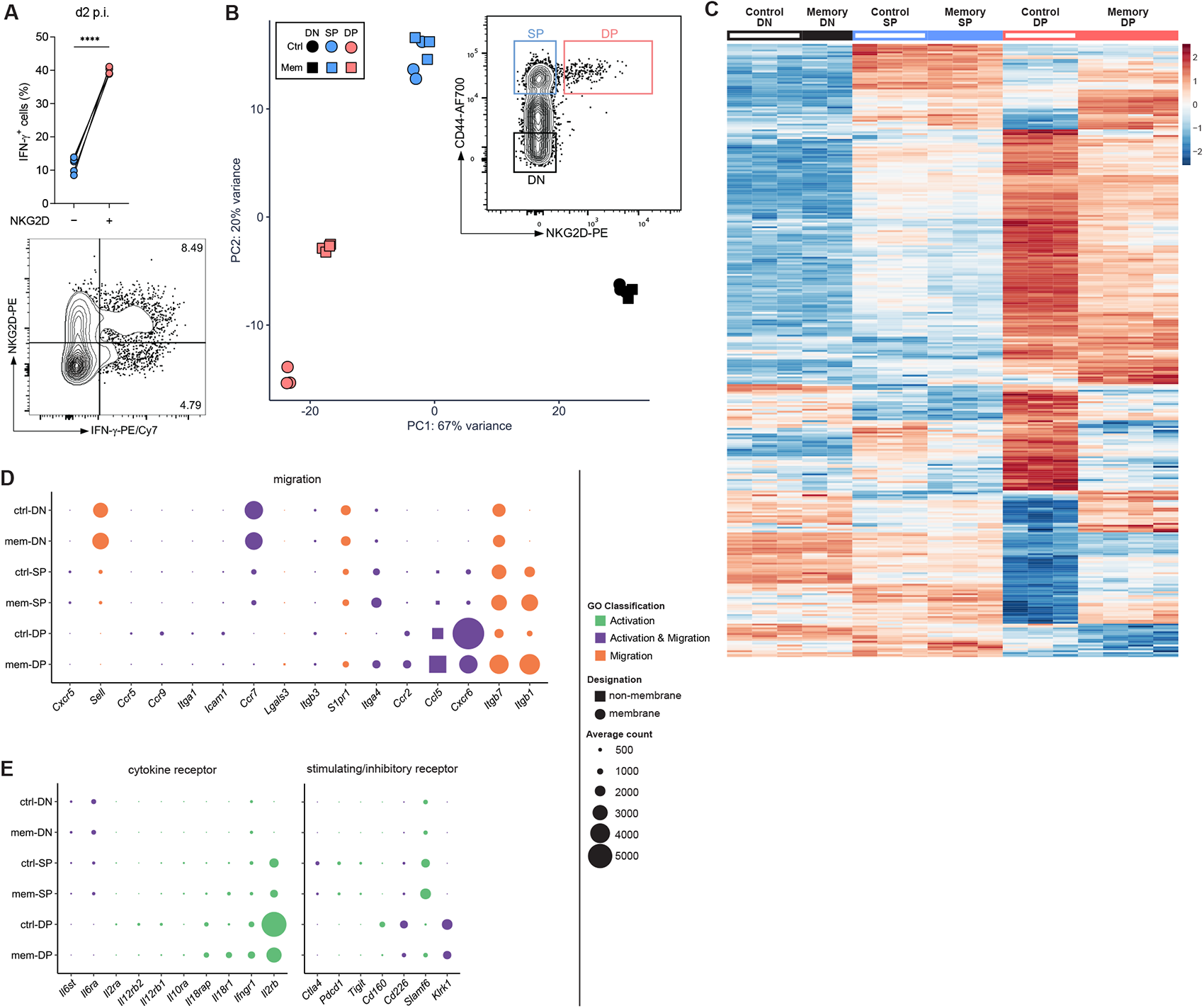
CD4⁺ T_IAM_ cells have a distinct transcriptional profile. **A**, IFN-γ response in NKG2D⁻ and NKG2D⁺ CD44^+^CD4^+^ T cells (top) and representative FACS plot gated on CD44^+^CD4⁺ T cells (bottom) obtained from lungs of LCMV memory mice challenged with *Lpn* (2 dpi; n = 5, paired t test). **B**–**E**, RNA-seq analysis of CD4⁺ T cells isolated from spleens of control (ctrl) and LCMV memory (mem) mice. **B**, Sorting strategy and principal component analysis of the resulting populations. **C**, Heatmap of genes in CD44^−^NKG2D^−^ (DN), CD44^hi^NKG2D^−^ (SP) and CD44^hi^NKG2D^+^ (DP) CD4^+^ T cells from LCMV memory and control mice (averaged normalized count from any population ≥ 10). **D**–**E**, differentially expressed genes (DEGs) relating to T cell migration or activation are highlighted, Balloon plots depicting average counts of the indicated genes for each cell population. DEGs are color-coded according to their GO classification. A curated list of migration- or cytokine receptor- and stimulatory/inhibitory receptor-related genes are shown.

Interestingly, control mice showed comparable numbers of splenic NKG2D^+^CD4^+^ T cells before Lpn infection but these were not recruited to the lung and could not mediate heterologous protection upon challenge (Figures S2C and S2D). To dissect how CD4^+^ T_IA_ cells are able to respond to heterologous challenges at distant sites, we performed transcriptional profiling of splenic NKG2D^+^CD4^+^CD44^+^ T cells from memory and control mice and compared them to NKG2D^−^CD4^+^CD44^+^ memory as well as CD4^+^CD44^−^ naïve T cells. NKG2D^+^CD4^+^CD44^+^ memory T cells from LCMV memory and control animals indeed formed distinct clusters, while CD4^+^CD44^−^ naïve and NKG2D^−^CD4^+^CD44^+^ memory cells from the two animal groups were transcriptionally very similar (Figures 2B, 2C, S3A, and S3B).

In line with a specific recruitment of memory CD4^+^ T_IA_ cells upon challenge, we observed differential expression of genes associated with cell migration when comparing NKG2D^+^CD4^+^CD44^+^ memory T cells from the two groups (e.g. *Itgb1*, *S1pr1*, *Itga4*) as well as genes upregulated in NKG2D^+^CD4^+^CD44^+^ compared to NKG2D^−^CD4^+^CD44^+^ memory or CD4^+^CD44^−^ naïve T cells (e.g. *Cxcr6*, *Ccr2*; Figures 2D, and S3B–S3D). Furthermore, comparison between NKG2D^+^ and NKG2D^−^ CD4^+^CD44^+^ memory or CD4^+^CD44^−^ naïve cells highlighted differential expression of a number of genes linked to T cell activation, including cytokine receptors (e.g. *Il2rb*, *Il18r1*) and stimulating/inhibitory receptors (e.g. *Cd226*, *Tigit*, *Pdcd1* and *Klrk1* encoding for NKG2D; Figures 2E, and S3A–S3D). Given that we found IFN-γ production by CD4^+^ T_IA_ cells to be TCR-independent (Figure 1G), we first tested whether IFN-γ secretion in memory CD4^+^ T cells could be induced by cytokines alone and focused on cytokines for which we observed differential expression of the receptors as well as those induced upon Lpn infection (IL-12, IL-18). While no single cytokine was able to induce IFN-γ production from memory CD4^+^ T cells, combination of IL-12 + IL-18, which are both induced upon Lpn infection (Spörri et al., 2008), and to a lesser degree IL-12 + IL-33, were able to stimulate IFN-γ secretion (Figures 3A and S4A). This is in line with a previous study reporting that IL-18 synergizes with IL-12 to produce IFN-γ in CD4^+^ T cells (Robinson et al., 1997; Guo et al., 2009). NKG2D^+^CD4^+^CD44^+^ T cells indeed showed higher expression of *Il12rb2* mRNA than NKG2D^−^CD4^+^CD44^+^ memory or CD4^+^CD44^−^ naïve T cells (Figure S4B). Furthermore, CD4^+^ T cells from memory mice showed a higher responsiveness to IL-12 as indicated by higher STAT4 phosphorylation upon *in vitro* cytokine stimulation as well as upon Lpn infection *in vivo* (Figures 3B and 3C). Memory CD4^+^ T cells also expressed higher levels of IL-18R (but not the IL-33 receptor ST2 encoded by *Il1rl1*; Figures S4B and S4C) and inhibition of the IL-18R signaling components p38 (SCIO469), JNK (SP60015), and AP-1 (SR 11302) reduced IFN-γ production by memory CD4^+^ T cells (Figures 3D and S4D). Finally, only CD4^+^ T cells from memory mice co-expressed the IL-18R on pSTAT4^+^ cells responding to IL-12 and were thus able to receive both signals necessary to induce IFN-γ production (Figure 3E). Despite the ability of IL-18 to induce IFN-γ production *in vitro*, blockade of IL-18R *in vivo* only resulted in a slight reduction of IFN-γ that did not reach significance and the IL-18 signal may thus be compensated by other stimulatory factors *in vivo* (Figure 3F). In contrast, IL-12 was essential for CD4^+^ T_IA_ cell activation *in vivo* as IL-12 blockade abolished the early IFN-γ response upon Lpn challenge (Figure 3F). Cytokines alone are thus sufficient to activate CD4^+^ T_IA_ cells during a heterologous challenge *in vivo* and CD4^+^ T_IA_ cells require a combination of two cytokines for TCR-independent activation, whereby IL-12 is essential.

**Figure 3.**
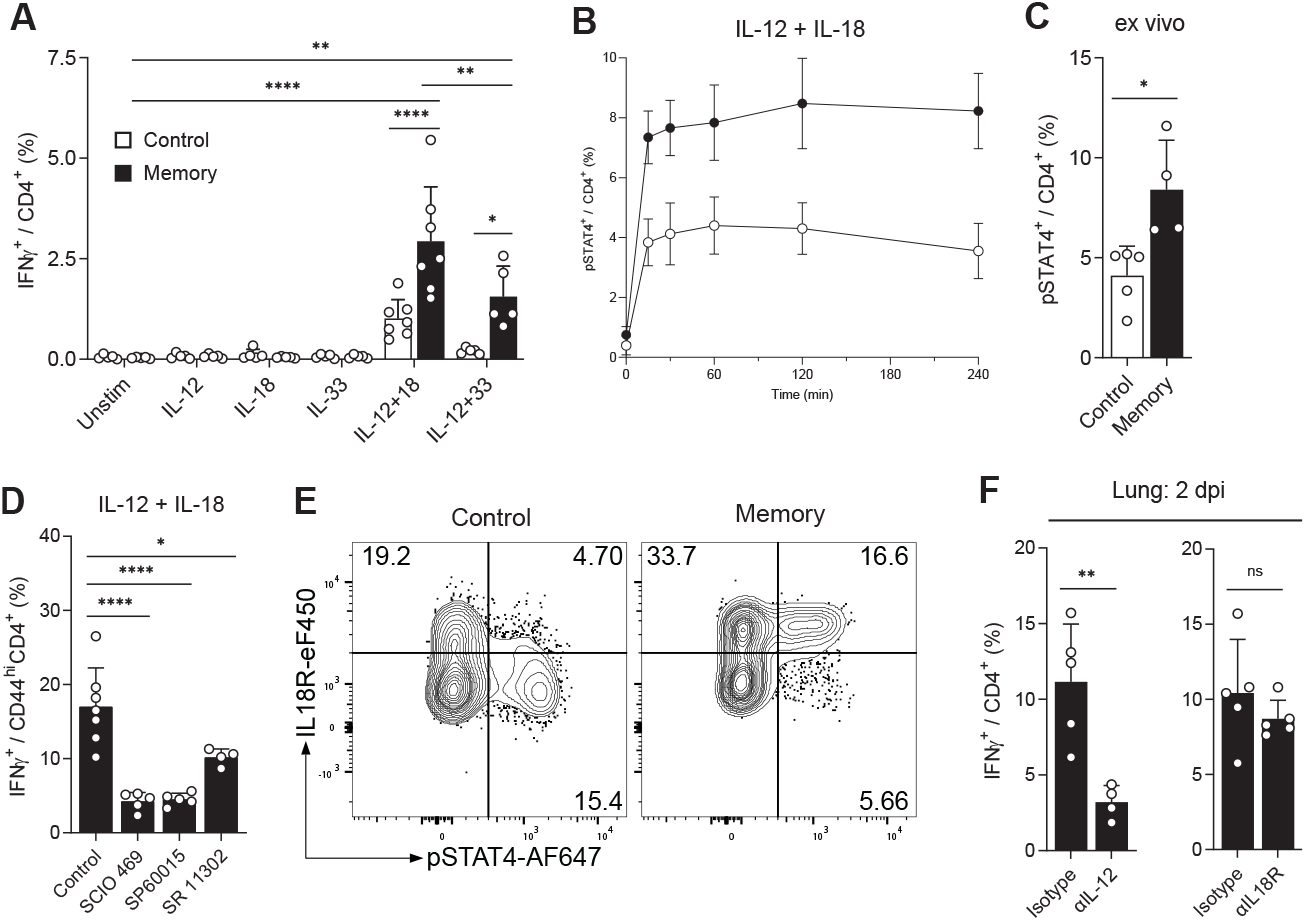
CD4⁺ T_IAM_ cells can be activated through cytokines. Splenic (A, B, D, and E) or lung (C and F) CD4⁺ T cells from control and LCMV-experienced memory mice were analyzed by flow cytometry. **A**, IFN-γ production after overnight stimulation with cytokines (n = 5–7). **B**–**C**, pSTAT4 levels after *in vitro* stimulation (B, n = 4), and pSTAT4 levels *ex vivo* 2 dpi. (C, n = 4–5). **D**, Cytokine stimulated cells as in (A) treated with inhibitors (n = 4–7). **E**, 2h stimulation with IL-12+IL-18, gated on CD44^+^CD4⁺ cells. **F**, *Ex vivo* analysis after intranasal treatment with neutralizing ɑIL-12 or blocking ɑIL18R (n = 4–5). Mean ± s.d., unpaired t test (C and F), Welch one-way (Dunnett T3; D) and ordinary two-way ANOVA (*Šídák*; A).

In line with the TCR-independent activation of CD4^+^ T_IA_ cells, 24h stimulation with cytokines did not result in a strong upregulation of co-inhibitory receptors, while stimulation of memory T cells through their TCR did induce such an upregulation (Figure S4E), as expected (Joller et al., 2011; Tietze et al., 2012). While in CD8^+^ T cells NKG2D can act as a co-stimulatory molecule and can also directly activate memory CD8^+^ T cells in a TCR-independent manner (Groh et al., 2001; Jamieson et al., 2002; Chu et al., 2013), NKG2D did not functionally contribute to the activation of memory CD4^+^ T_IA_ cells as stimulation with blocking or agonistic anti-NKG2D antibodies had no effect on the IFN-*γ* response (Figure S4F). In line with these results, NKG2D expression is a poor predictor of the magnitude of the early, TCR-independent IFN-*γ* response (Figure S4G), despite the fact that NKG2D^+^CD4^+^CD44^+^ T cells were most potent producers of IFN-*γ* upon cytokine stimulation (Figure S4H). Additionally, it is important to note that not all CD4^+^CD44^+^ memory T cells have this responsiveness to cytokine stimulation, as only a small fraction of memory CD4^+^ T cells was able to produce IFN-*γ* (Figure S4H). NKG2D thus acts as a marker rather than a functionally relevant receptor of CD4^+^ T_IA_ cells, which are activated in a TCR- and NKG2D-independent manner by cytokine alone whereby IL-12 is essential for their activation *in vivo*.

### Recruitment of CD4^+^ T_IA_ cells

To determine how the recruitment of CD4^+^ T_IA_ cells to the site of challenge is regulated, we further investigated molecules associated with T cell migration that were differentially expressed in NKG2D^+^CD4^+^CD44^+^ T cells from memory vs. control mice and in NKG2D^+^ vs. NKG2D^−^ CD4^+^CD44^+^ memory or CD4^+^CD44^−^ naïve T cells (Figures 2D, S3D and S5A). In line with the transcriptional data, we could observe a very high expression of the chemokine receptor CXCR6 on NKG2D^+^CD44^+^CD4^+^ T cells (Figure S5A) and IFN-*γ*^+^ CD4^+^ T_IA_ cells (Figure S5B). Indeed, CXCR6 together with IL-18R was an even better marker for IFN-γ expression than NKG2D (Figure 4A), and additionally served as an excellent predictor of the magnitude of the IFN-γ response (Figure 4B). Nevertheless, CXCR6 was highly expressed in CD4^+^ T_IA_ cells from memory mice but also in NKG2D^+^CD4^+^CD44^+^ T cells from control mice, which could be stimulated to produce IFN-γ *in vitr*o, but were not recruited to the site of infection *in vivo* (Figures S2C, S2D and S4H), hinting towards alternative recruitment mechanisms. Indeed, blocking of the CXCR6 ligand CXCL16 did not alter CD4^+^ T_IA_ cell recruitment or IFN-γ production and CXCR6 is thus not essential for migration of CD4^+^ T_IA_ cells to the site of infection (Figure S5C).

**Figure 4.**
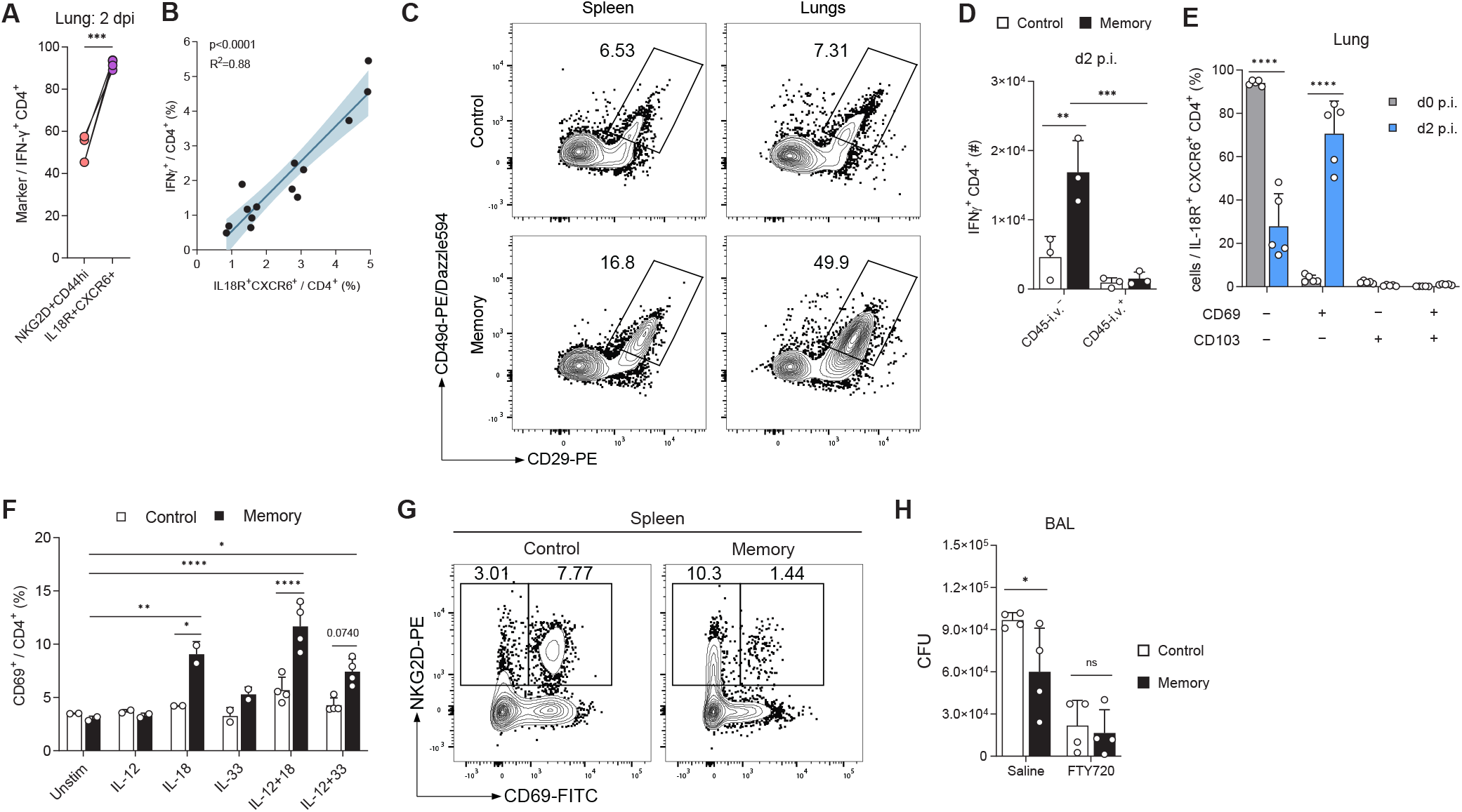
CD4⁺ T_IAM_ cells have enhanced migratory capabilities. **A**, *Ex vivo* expression of indicated markers upon Lpn infection (n = 5). **B**, Linear regression of splenic control and memory CD4⁺ T cells stained *ex vivo* (x-axis) and IFN-γ production upon overnight IL-12+IL-18 stimulation (y-axis). 95% confidence interval is indicated (n = 14). **C**, Representative FACS plot of integrin expression of CD4^+^ T cells 2 dpi. **D**, Mice injected with ɑCD45 antibody i.v. prior to sacrifice (n = 3). **E**, Indicated marker expression among T_IAM_ cells (n=5). **F**, Isolated splenic CD4+ T cells were incubated with indicated cytokines overnight (n=2–4). **G**, Representative FACS plot of CD44^+^CD4^+^ T cells pre-Lpn infection. H, Lpn titers 3 dpi. in mice treated with FTY720 or saline i.p. (n = 4). Mean ± s.d., paired t test (A) and ordinary two-way ANOVA (*Šídák*; D, E, F, and H).

Besides CXCR6, CD4^+^ T_IA_ cells display high expression of the integrin VLA-4 (constituted of CD49d and CD29 encoded by *Itga4* and *Itgb1*; Figures 2D, 4C, S5A, and S5D), which plays an important role in lymphocyte homing and tissue entry (Elices et al., 1990). To test whether CD4^+^ T_IA_ cells preferentially enter the lung tissue upon Lpn challenge, we injected a fluorescently labelled anti-CD45 antibody intravenously shortly before sacrifice to distinguish cells in the vasculature and tissue of the lungs. Lpn-challenged memory mice indeed harbored more CD45-i.v. negative IFN-γ^+^ T cells that had entered the lung tissue than control mice (Figures 4D and S5E). Absence of CD45-i.v. labelling could be the consequence of tissue entry upon recruitment or indicate a previously established niche of tissue resident memory CD4^+^ T cells in the lung. To investigate whether CD4^+^ T_IA_ cells exhibit tissue residency features, we analyzed their expression of the tissue residency markers CD69 and CD103 (Y.-T. Lee et al., 2011; Turner et al., 2014). Analysis of CD69 and CD103 expression in the lung pre- and post-Lpn challenge revealed a lack of tissue residency marker expression among CD4^+^ T_IA_ cells prior to the Lpn infection (Figure 4E). At day 2 post Lpn infection, CD4^+^ T_IA_ cells were still negative for CD103, confirming that they do not represent a tissue resident memory population. Nevertheless, the majority of CD4^+^ T_IA_ cells expressed CD69 after Lpn challenge. Upregulation of CD69 has been reported to occur upon tissue entry and it is speculated that local factors contribute to this induction (Cibrián and Sánchez-Madrid, 2017). Because of the cytokine-mediated activation of CD4^+^ T_IA_ cells (Figures 3A and 3F), we wondered whether these cytokines could induce CD69 expression. Indeed, LCMV-experienced splenic CD4^+^ T cells, but not naïve controls, showed an upregulation of CD69 upon overnight stimulation with IL-18 alone or in combination with IL-12 and to a lesser degree by stimulation with IL-12+IL-33 (Figure 4F). These findings indicate that CD4^+^ T_IA_ cells can preferentially enter peripheral tissues where they then upregulate CD69 and are retained upon cytokine activation.

Interestingly, both control and memory NKG2D^+^CD4^+^CD44^+^ T cells expressed elevated levels of S1PR1 (Figures 2D, S5F, and S5G), a G-protein-coupled receptor required for lymphocyte egress from lymphoid organs (Matloubian et al., 2004). However, in contrast to NKG2D^+^CD4^+^CD44^+^ T cells from control mice, prior to challenge, splenic CD4^+^ T_IA_ cells were negative for CD69 (Figure 4G), which is known to promote T cell retention in the spleen and acts as a negative regulator of S1PR1 (Shiow et al., 2006). Absence of CD69 together with a high expression of S1PR1 thus equip CD4^+^ T_IA_ cells with a superior ability to exit the spleen upon challenge (Figure S5G). Finally, complete blockade of T cell egress from secondary lymphoid organs using fingolimod abolished the early heterologous protection we observed (Figure 4H), highlighting the importance of CD4^+^ T_IA_ cell migration for their protective function.

### CD4 T_IA_ cells promote autoimmunity

T cell migration to and cytokine production at tissue sites are not only essential in immunity to infections but also play an important role in autoimmune disorders. To determine whether CD4^+^ T_IA_ cells may contribute to the etiology of autoimmunity, we first assessed whether they are present in autoimmune settings. EAE is a well-established model for multiple sclerosis and can be induced by active immunization with CNS antigens or by adoptive transfer of activated T cells specific for CNS antigen such as 2D2 cells, which recognize a peptide derived from myelin (Bettelli et al., 2003; Jäger et al., 2009). Using the adoptive transfer EAE model, we found that CD4^+^ T_IA_ cell markers NKG2D^+^ or CXCR6^+^IL-18R^+^ were not only highly enriched on CNS antigen-specific 2D2 cells but also on non-specific endogenous T cells, supporting the notion that CD4^+^ T_IA_ cells are also recruited to sites of autoimmune inflammation in an antigen-independent manner (Figures 5A, 5B, S6A, and S6B). This is in line with reports that found increased frequencies of NKG2D^+^ CD4^+^ or CXCR6^+^ CD4^+^ T cells at the site of autoimmune inflammation in patients suffering from multiple sclerosis, rheumatoid arthritis or systemic lupus erythematosus (Groh et al., 2003; van der Voort et al., 2005; Yang et al., 2009; Ruck et al., 2013; Beltrán et al., 2019; Hiltensperger et al., 2021). Mirroring their effector function upon infectious challenge, NKG2D^+^ and CXCR6^+^IL-18R^+^ CD4^+^ T cells were higher producers of IFN-*γ*in active and passive models of EAE (Figures 5C, 5D, S6C, and S6D). Importantly, CNS-infiltrating endogenous CD4^+^ T cells could be potently activated to produce high amounts of IFN-γ when stimulated with IL-12+IL-18 alone (Figure 5E), confirming that true CD4^+^ TIA cells capable of massive cytokine release in the absence of a TCR signal are indeed present at the site of autoimmune response.

**Figure 5.**
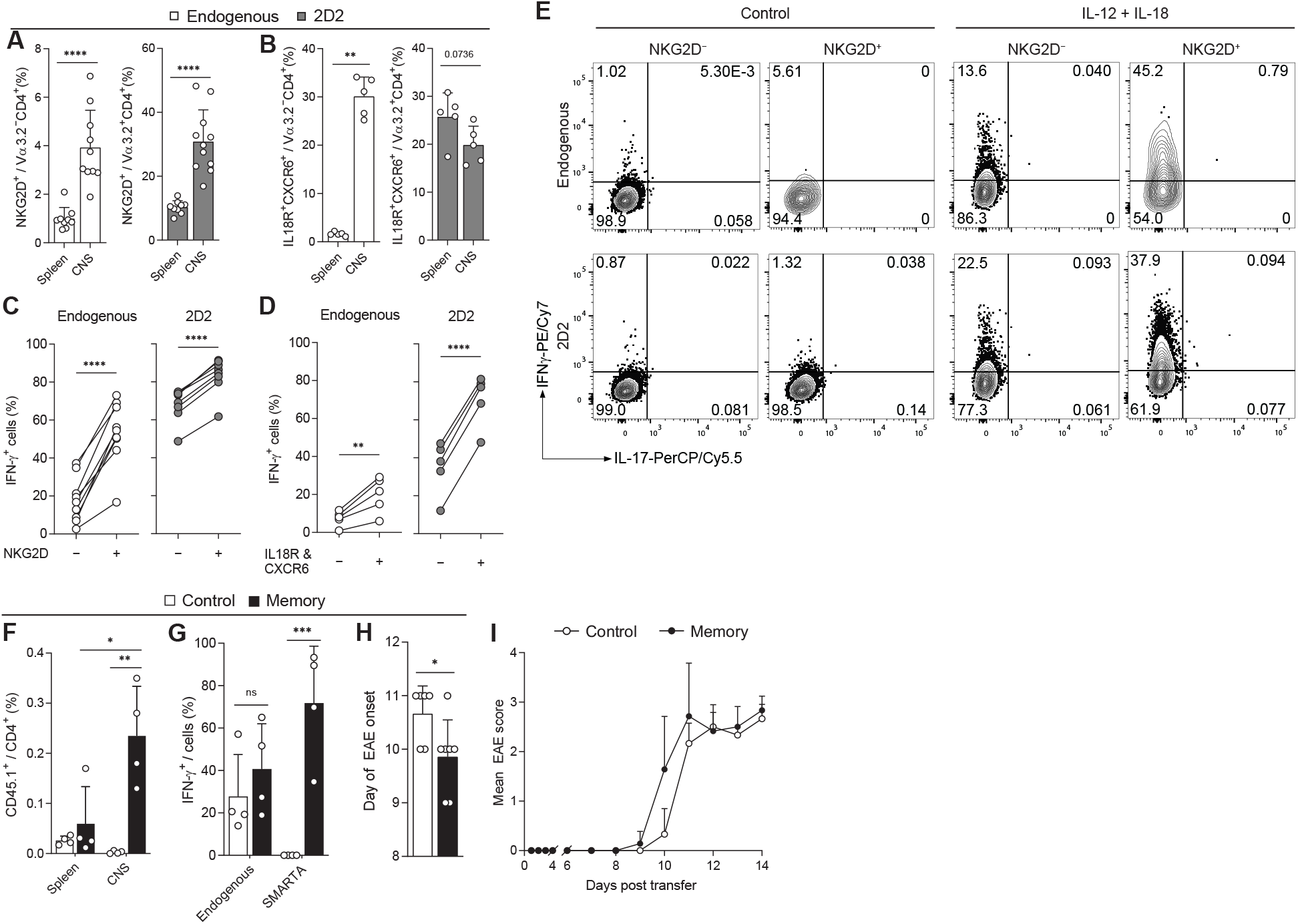
CD4^+^ T_IAM_ cells promote CNS inflammation in EAE. For induction of EAE, Th1 polarized 2D2 cells were transferred into WT mice and identified by Vα3.2 expression (A–E). **A**–**B**, Expression of NKG2D (A, n = 9–11) and IL-18R and CXCR6 (B, n = 5) in CD4^+^ T cells was determined by flow cytometry at the peak of EAE. **C**–**D**, IFN-g response among NKG2D⁻ or NKG2D^+^ (C, n = 8–9) and IL-18R^−^CXCR6^−^ or IL-18R^+^CXCR6^+^ (D, n = 5) CD4^+^ T cells from the CNS at the peak of disease. **E**, Cells isolated from the CNS were stimulated with IL-12+IL-18 or left untreated overnight and analyzed for IFN-g production, plots are gated on CD4^+^ T cells. **F**–**G**, MOG_35-55_/CFA-immunized C57Bl/6 mice received 2×10^6^ naïve or memory SMARTA CD4^+^ T cells (CD45.1^+^). SMARTA frequency (F) and IFN-g response in the CNS (G) (n = 4). **H**–**I**, 5×10^5^ 2D2 CD4^+^ T cells and 5×10^5^ naïve or memory SMARTA cells were co-transferred into *Rag1*^−/−^ mice immunized with MOG_35-55_/CFA and monitored for disease (n = 6–7). Mean ± s.d., Mann-Whitney test (A), unpaired (B, H) and paired t-test (C and D), two-way ANOVA (*Šídák*; F and G).

Next, we tested our hypothesis that virus-experienced memory CD4^+^ T_IA_ cells are preferentially recruited to sites of autoimmune inflammation. To this end, we transferred naïve or memory Smarta T cells, which do not recognize CNS antigens, into EAE recipient mice. In line with our hypothesis, we found that memory but not naïve Smarta T cells infiltrated the CNS of EAE mice (Figures 5F and S6E) and produced high amounts of IFN-*γ* even in the absence of their cognate antigen (Figures 5G and S6E). Finally, co-transfer of LCMV-specific memory but not naïve Smarta cells together with MOG-specific 2D2 cells accelerated EAE disease onset (Figures S6F, 5H, and 5I), confirming that CD4^+^ T_IA_ cells can contribute to the development of autoimmunity. Like in heterologous infectious challenges, CD4^+^ T_IA_ cells are thus preferentially recruited to the site of autoimmune inflammation, where they can be activated in a TCR-independent manner to produce IFN-*γ* and contribute to autoimmune disease.

## Discussion

The capability to generate memory cells upon pathogen encounter is one of the greatest advantages of the vertebrate immune system. Memory T cells mount an accelerated and augmented response upon re-encounter of their cognate antigen resulting in enhanced pathogen control. Here, we show that memory CD4^+^ T cells generated in response to a viral infection are also capable of mounting an early IFN-*γ* response in unrelated heterologous challenges. This rapid and antigen-independent response exerted by CD4^+^ T_IA_ consequently modulates disease susceptibility in infectious and autoimmune settings.

Following heterologous bacterial challenge, CD4^+^ T_IA_ cells reduced the bacterial burden in a TCR-independent manner. Importantly, CD4^+^ T_IA_ cells alone were able to confer this protection, confirming their functional relevance *in vivo*. CD4^+^ T_IA_ cells established following LCMV infection have a distinct transcriptional profile with upregulated expression of cytokine and chemokine receptors such as IL-18R and CXCR6. Similar to memory CD8^+^ T cells (Freeman et al., 2012), CD4^+^ T_IA_ cells can be stimulated *in vitro* to produce IFN-γ by the cytokine combination IL-12 + IL-18, both of which are present during Lpn infection (Spörri et al., 2008). *In vivo*, IL-12 was essential for the CD4^+^ T_IA_ cell response. In contrast, we did not see a reduction of IFN-γ production by blocking signals through the IL-18R. This fits with our observation that IL-33 in combination with IL-12 was also able to stimulate IFN-γ secretion, although to a lesser degree. While IL-18 and IL-33 are both IL-1 family cytokines, IL-1*β* itself could not synergize with IL-12 to activate CD4^+^ T_IA_ cells. Whether this is due to the nature of the signal of each specific receptor or the ability of CD4^+^ T_IA_ cells to sense these cytokines remains to be determined. Furthermore, the possibility that the second signal could also be delivered through costimulatory molecules, such as e.g. LPS (Nogai et al., 2005), rather than from cytokines, has not been excluded. Thus, while IL-12 is essential for TCR-independent re-activation of CD4^+^ T_IA_ cells, a certain redundancy exists for the second signal necessary to evoke this heterologous response.

Interestingly, these cytokine combinations could also induce IFN-γ production in CD44^+^ T cells from control animals that to a certain degree resemble CD4^+^ T_IA_ cells and were likely generated in response to commensals or dietary antigens. However, despite their responsiveness to the cytokine stimulation, CD44^+^ T cells from control animals did not have the capacity to migrate to the site of heterologous challenge, which was crucial for the protective effect of CD4^+^ T_IA_ cells in memory mice. Although CXCR6 (together with IL-18R expression) serves as a good predictor of the IFN-*γ* response, it does not functionally contribute to CD4^+^ T_IA_ cell recruitment. This is in line with previous reports that found CXCR6 to be dispensable for CD4^+^ T cell migration to the site of inflammation (L.N. Lee et al., 2011; Hiltensperger et al., 2021). Interestingly, a recent study revealed that the intestine forms a reservoir for Th17 cells from which pathogenic cells then disseminate via the spleen to sites of autoimmune inflammation (Schnell et al., 2021). The reduction of CD4^+^ T_IA_ cell numbers in spleen we observed following heterologous challenge suggests that the spleen may similarly serve as a kind of reservoir for Th1 CD4^+^ T_IA_ cells that can then be rapidly recruited to peripheral sites upon challenge. CD4^+^ T_IA_ cells are thus set apart from CD4^+^ memory T cells in control mice by their ability to be recruited to the site of heterologous challenge where they then encounter a cytokine environment that allows for their TCR-independent activation and modulation of the immune response.

CXCR6^+^CD4^+^ T cells recently received much attention as prominent producers of cytokines and highly pathogenic effector cells during autoimmunity (Hou et al., 2019; Hiltensperger et al., 2021; Schnell et al., 2021). While these studies focus on antigen-specific cells, we show here that CD4^+^ T_IA_ cells that are activated in a TCR-independent manner contribute to this pool of pathogenic cells and drive autoimmune disorders, as we have seen for EAE. Viral infections have long been discussed as triggers for many autoimmune diseases and the rise in patients presenting with autoimmune symptoms following SARS-CoV-2 infections has strengthened the link between viruses and autoimmunity (Toscano et al., 2020; Verdoni et al., 2020). However, except for some rare instances (Lanz et al., 2022), no causal link could be established between the virus and the autoimmune disease (Getts et al., 2013). Our study suggests that, in addition to rare cases in which cross-reactive cells are established following viral infections (Lang et al., 2002), CD4^+^ T_IA_ cells generated as part of the virus-specific memory response are recruited to the site of autoimmune inflammation where they are activated to produce IFN-*γ*. Collectively, our findings demonstrate that, upon challenge, memory CD4^+^ T cells are recruited to inflamed tissue sites and contribute to the local inflammatory response in an antigen-independent manner, thus favoring pathogen control but also promoting autoimmune pathology. This early, innate-like and antigen-independent nature of the T_IA_ cell-response outlined in our study has highlighted an additional functionality of T cells to exert effector functions beyond the classical antigen-driven, delayed adaptive immune response.

## Materials and Methods

### Mice

C57BL/6 (B6) mice were purchased from Janvier Labs, and Rag1-KO mice were purchased from Jackson laboratories. Congenic Ly5.1 and Thy1.1, Smarta (Oxenius et al., 1998), 2D2 (Bettelli et al., 2003), and *Rag2*^−/−^γc^−/−^(Song et al., 2010) mice have been described previously. All animals were bred and housed in SPF and OHB facilities at LASC Zürich, Switzerland or in the CAM in Munich, Germany. All experiments were performed in accordance with institutional policies and regulations of the relevant animal welfare acts and have been reviewed and approved by the Cantonal veterinary office or by the local animal ethics committee of the state of Bavaria (Regierung von Oberbayern) in accordance with European guidelines.

### Viruses, bacteria, and infections

The LCMV WE strain was propagated on L929 cells and titrated on MC57G cells and animals were infected i.v. with 200 FFU to induce an acute infection. Vaccinia Virus was propagated on BSC40 cells and mice were infected with 10^6^ PFU i.p.. 40–100 days after the primary infection, mice were intranasally infected with 3×10^6^ CFU *L. pneumphila* JR32 FlaA^−^ (Lpn) (Weber et al., 2012), grown on charcoal yeast agar plates. At sacrifice, animals were perfused with PBS and bronchioalveolar lavage (BAL), lung, and spleen were collected. For determination of bacterial titers, BAL and lungs from infected mice were collected, lungs were lysed using a Qiagen TissueLyser II and samples were plated on charcoal yeast agar plates and grown for 3 days at 37 °C.

For the blockade of IL-18R or IL-12, Lpn was co-administered i.n. together with 20 µg anti-IL18R antibody (clone 112624; R&D Systems) or 60 µg of anti-IL-12p40 antibody (clone C17.8; BioLegend). IFN-γ was neutralized by giving 100 µg anti-IFN-γ antibody (clone XMG1.2; BioLegend) i.n. on day 0 and 1 of the Lpn infection. To block the ligand of CXCR6, 100 µg of anti-CXCL16 antibody (clone 142417; R&D Systems) was injected i.v. FTY720 was injected daily at 1 mg/kg i.p until sacrifice starting one day before Lpn infection.

### Adoptive cell transfers

For adoptive transfers and *in vitro* assays, CD4^+^ T cells were purified using MojoSort Mouse CD4 Nanobeads (BioLegend). For transfer into *Rag2*^−/−^g*c*^−/−^ mice, purified CD4^+^ T cells obtained from spleen were additionally sorted for CD4 expression. To generate memory Ly5.1 Smarta cells, 10^4^ cells were adoptively transferred i.v. into B6 recipient mice one day prior to LCMV infection (200 FFU LCMV WE i.v.). Memory cells were obtained from spleens using the MojoSort Mouse CD45.1 selection kit (BioLegend) >40 days post infection.

To induce EAE by adoptive cell transfer, naïve CD4^+^ T cells were isolated from spleen and lymph nodes of 2D2 mice. To prepare a single-cell suspension, spleens and lymph nodes were mashed and passed through a 70 μm cell strainer. After erythrocyte lysis, naïve CD4+ T cells were purified using the naïve CD4^+^ T cell isolation Kit (Miltenyi Biotec). Naïve T cells were cultured at a concentration of 1.5–2×10^6^/ml in complete RPMI 1640 medium (supplemented with 10% heat-inactivated FBS, 1% penicillin-streptomycin, 10 mM HEPES, 2 mM L-glutamine, 1% non-essential amino acids, 1 mM sodium pyruvate and 50 μM *β*-Mercaptoethanol) in the presence of 7.5–10×10^6^/ml irradiated (35 Gy) splenocytes and 2.5 µg/ml soluble anti-CD3 antibody (clone 145-2C11, BioXCell). Th1 cells were generated by addition of IL-12 at a concentration of 10 ng/ml and anti-IL-4 antibody (clone 11B11, BioXCell) at a concentration of 10 µg/ml into the culture. For the generation of Th17 cells, naïve T cells were cultured with IL-6 at a concentration of 30 ng/ml, TGF-ß at a concentration of 3 ng/ml, IL-1ß at a concentration of 20 ng/ml, and anti-IFN-γ (clone XMG1.2, BioXCell) and anti-IL-4 Ab (clone 11B11, BioXCell) at a concentration of 10 µg/ml. After 48h, Th1 cells and Th17 cells were split with medium containing 10 ng/ml of IL-2 and medium containing 10 ng/ml of IL-23, respectively. All cytokines were purchased from BioLegend except IL-23 (Miltenyi Biotec). The different T cell subsets were analyzed for cytokine production after 4 days. After 5–8 days, cells were restimulated at a concentration of 2×10^6^/ml for 48h in the presence of plate-bound anti-CD3 and anti-CD28 (clone PV-1; BioXCell) antibodies both at 2 µg/ml in fresh medium without any cytokines. 2–4×10^6^ cytokine-producing cells were injected i.p. into B6 recipients.

For experiments using SMARTA cells in the setting of EAE, either 2×10^6^ naïve or memory SMARTA cells were injected i.v. into C57BL/6 mice after MOG_35-55_/CFA immunization or 5×10^5^ naïve or memory SMARTA cells (together with 5×10^5^ 2D2 CD4^+^ T cells) were injected i.v. into RAG1 KO mice with MOG_35-55_/CFA immunization performed on the following day.

### Flow cytometry

FACS stainings were performed on single-cell suspensions from spleen, lung, BAL, and CNS. Where indicated, mice received 2 µg anti-CD45.2 APC antibody (clone 104) in 200 µl PBS intravenously before sacrifice. 3 min following administration they were sacrificed by anesthesia (Isoflurane) and cervical dislocation. The spleen samples were prepared by mechanical disruption in RPMI 1640 medium supplemented with 10% FCS, penicillin (100IU/ml) and 1% L-glutamine. Lungs were enzymatically digested with collagenase D (Gibco) and DNase I (VWR) for 30 min, and immune cells isolated using a 30% Percoll (GE Healthcare) gradient. Red blood cells were lysed with ACK buffer (155 mM NH_4_Cl, 10mM KHCO_3_, 0.1 mM Na_2_EDTA, pH: 7.4) for 3 min. For Lpn re-stimulation, cells were stimulated with Lpn-extract at 37 °C in 10% CO_2_ for 6h before staining. When staining for intracellular cytokines, cells were incubated with Brefeldin A (BioLegend) for 4h at 37 °C in 10% CO_2_ prior to staining. For surface stainings, antibodies were incubated for 20–30 minutes at RT in PBS. The Zombie NIR fixable dye (Biolegend) was used to exclude dead cells and debris. For intracellular cytokine staining, cells were permeabilized using the Cytofix/Cytoperm kit (BD Biosciences) for 5–8 minutes at RT, followed by antibody incubation for 20–30 minutes at RT. To stain for phosphorylated STAT4, cells were incubated for 12 min at 37 °C with PFA (4%), and upon washing, fixed with 90% methanol for 30 min on ice. After fixation with methanol, cells were stained for 45 min at RT.

For EAE experiments, cells were stimulated with PMA (50 ng/ml, Sigma-Aldrich) and ionomycin (500 ng/mL, Sigma-Aldrich) in the presence of monensin (0.7 µL/mL, GolgiStop; BD Biosciences) at 37 °C in 5% CO2 for 3.5 before staining. For surface stainings, antibodies were incubated for 20–30 min at 4 °C in PBS + 2% FBS. For intracellular cytokine staining, cells were fixed for 30 min at 4 °C with 0.4% paraformaldehyde (Merck KGaA) and permeabilized with PBS containing 2% FBS and 0.1% saponin (Sigma-Aldrich), followed by antibody incubation for 30 min at 4 °C. The Zombie UV fixable viability kit (Biolegend) was used to exclude dead cells and debris.

The following antibodies were used for flow cytometry: αCD45-eF450 (30-F11), αIL-17A-APC (eBio17B7), αIL-18Ra-PE (P3TUNYA), streptavidin-APC, rat IgG1 κ-PE-Cy7 isotype control (eBRG1), rat IgG2α κ-PE isotype control (eBR2a) were all purchased from eBioscience. αCD11b-APC or AF700 (M1/70), αCD4-BV605 (RM4-5), αIFNγ-PE-Cy7 (XMG1.2), αB220-PerCP-Cy5.5 (RA3-6B2), αVα3.2-FITC (RR3-16), αIL-10-PE (JES5-16E3), αCD19-PerCP-Cy5.5 (1D3), αCD45.1-FITC (A20), αCD45.2-PerCP-Cy5.5 (104), biotin αCXCR6 (SA051D1), αNKG2D-PE (CX5), αIL-17A-PerCP-Cy5.5 (TC11-18H10.1), rat IgG2α κ-PerCP-Cy5.5 or APC isotype control (RTK2758), rat IgG2b κ-PE isotype control (RTK4530), rat IgG1 κ-PE isotype control (RTK2071) were all purchased from Biolegend. FACSAria III was used for sorting of cells. Data was acquired on a BD LSR Fortessa, BD FACS CantoII, BD FACSverse, or BD FACSymphony A5 analyser (BD Bioscience) and analysed using Flowjo software (TreeStar).

### CyTOF

Single cell suspensions obtained from lung and spleens of control and LCMV-memory mice challenged with Lpn were incubated with Brefeldin A, labelled, and prepared for cytometry by time of flight (CyTOF) according to manufacturer’s instructions and as previously described (Johansson et al., 2021). In brief, all samples were stained with cisplatin (Fluidigm #20164; used to determine live cells), fixed with Fluidigm MaxPar® Fix I Buffer (Fluidigm #201067) and barcoded using Fluidigm Cell-ID 20-Plex Pd Barcoding Kit (Fluidigm #201060). Subsequently, all barcoded lung and spleen samples were pooled into one lung and one spleen sample mix, respectively, and stained with the cocktail of monoisotope-labelled antibodies listed in Supplementary Table 1. Of note, the antibodies obtained from Fluidigm were purchased already labeled by the vendor, whereas the antibodies obtained from Biolegend were labelled in house using the specific Maxpar® antibody labelling kits from Fluidigm. Following antibody staining, the pools of lung and spleen samples were washed with MaxPar® Cell Staining Buffer (Fluidigm #201068), resuspended with Cell-ID™ Intercalator-Ir solution (Fluidigm #201192B; used to assess single cell events) and left overnight at 4 °C. Next day, cells were washed again MaxPar® Cell Staining Buffer, resuspended in MaxPar® water (Fluidigm #201069), pelleted and stored dry until acquisition. Immediately before data acquisition, the lung and spleen cell pellets were adjusted to 1×10^6^ cells/ml in MaxPar® water containing 10% EQ Four Element Calibration Beads (Fluidigm #201078; used to normalize data for signal variation occurring over acquisition time). Data acquisition was performed using a Fluidigm (Helios™) mass cytometer. Fcs data files were normalized with a software tool provided by Fluidigm, and deconvoluted according to barcodes and analyzed in FlowJo.

### In vitro stimulation

CD4^+^ T cells were isolated using MojoSort Mouse CD4 T cell isolation kit (BioLegend). Upon isolation, cells were incubated with the indicated cytokines (10–100 ng/ml) for 12–16h overnight. Inhibition of the MyD88-pathway was investigated by co-incubating 1 µM SCIO 469 (TOCRIS), 5 µM SP600125 (Sigma), and 20 µM SR 11302 (TOCRIS) together with the IL-12 + IL-18 stimulation overnight. For *in vitro* antibody stimulation, flat bottom plates were coated with anti-CD3 (2 µg/ml, clone 145-2C11; BioXCell) and anti-CD28 (2 µg/ml, clone PV-1; BioXCell) or anti-NKG2D (10 µg/ml, clone CX5 or A10; BioLegend). Recombinant cytokines were purchased from Biolegend.

### RNA sequencing

Cells isolated from spleen were sorted into 96-well plates (500 cells / well) by using a single cell mask. For RNA isolation, the Smart-Seq2 protocol was applied as described in (Schorer et al., 2020). Briefly, Agencourt RNAClean XP paramagnetic beads (Beckman Coulter) were used in combination with a DynaMag-96 side skirted magnet (Thermo Fisher). cDNA was generated with SuperScript II Reverse Transcriptase Kit (Thermo Fisher) and amplified with HiFi HotStart PCR Mix (KAPA Biosystems). For DNA clean-up Agencourt AMPure XP beads were used (Beckman Coulter) as above. Nextera XT DNA sample preparation and index kits (Illumina) were used for preparation of libraries that were sequenced by the Functional Genomics Center Zurich (Zurich, Switzerland).

### Analysis of RNA sequencing data

Raw sequencing files were aligned to the mouse genome (GRCm38) with HiSat2 (Wen, 2017) (version 2.2.1) following quality control with FastQC (https://www.bioinformatics.babraham.ac.uk/projects/fastqc/; version 0.11.9). Count tables were generated with featureCounts (Liao et al., 2014) (version 1.22.2) using the options ‘-t exon -g gene_id’ and the GTF file of the GRCm38 build (version 101) as reference. Data were analyzed using R (version 4.0.2). DESeq2 (Love et al., 2014) (version 1.28.1) was used for normalization of counts and principal component analysis (PCA). Differentially expressed genes (DEGs) were defined by adjusted p-value <0.05 and fold differences >2. Genes that had lower counts than 250 in all group averages were disregarded. Gene information together with gene ontology (GO) entries were obtained with Ensembl (version 101) using the biomaRt package (Durinck et al., 2005; Durinck et al., 2009) (version 2.46.3). GO entries were used to group genes into the categories “activation” (GO entries: “activation” OR “immune response” OR “cytokine” OR “positive regulation of cell cycle”), “migration” (GO entries: “taxis” OR “migration” OR “chemokine” OR “cell adhesion”), “activation & migration” (activation AND migration), “transcription factor” (GO entry: “DNA-binding transcription factor activity”), “regulator of transcription factor” (GO entry: “regulation of DNA-binding transcription factor activity”). Furthermore, genes were distinguished by their association with cell membrane/being at the cell surface (“integral component of membrane” OR “cell surface” OR “anchored component of plasma membrane”). To plot the results, the packages ggplot2(Wickham, 2009) (version 3.3.3), pheatmap (version 1.0.12), UpSetR(Conway et al., 2017) (version 1.4.0), VennDiagaram (version 1.6.20) were used.

### Quantitative RT-PCR

CD4^+^ T cells were isolated from the spleen using MojoSort Mouse CD4 T cell isolation kit (BioLegend) and sorted according to their expression of CD44 and NKG2D. After the sort, cells were taken up in Buffer RLT (Qiagen) and stored at −20 °C. For RNA extraction, RNeasy Kit (Qiagen) was used by following the manufacturer’s instructions. Following the extraction, cDNA was created using High-Capacity cDNA Reverse Transcription Kit (Applied Biosystems). For measurement of relative gene expression, real-time quantitative PCR (RT-qPCR) was performed using TaqMan Fast Advanced Master Mix (Applied Biosystems) and the following primers which were all purchased from Applied Biosystems: *Actb* (Mm00607939_s1), *Il12rb2* (Mm01183807_m1), *Il18r1* (Mm00515178_m1), *Il1rl1* (Mm00516117_m1), *S1pr1* (Mm02619656_s1). All measurements were acquired on Bio-Rad CFX384 Touch Real-Time PCR Detection System and cycle threshold values were obtained through the CFX Maestro software (Bio-Rad).

### EAE induction and scoring

B6 mice were immunized with 100–200 µg MOG_35-55_ (BioTrend) emulsified in CFA (Difco Laboratories) containing 5 mg/ml Mycobacterium tuberculosis (Difco Laboratories). Additionally, they received 150ng PT (List laboratories) on day 0 and 2 after immunization. To induce EAE in RAG1 KO mice, CD4^+^ T cells were isolated from spleen and lymph nodes of 2D2 mice. To prepare a single-cell suspension spleens and lymph nodes were mashed and passed through a 70-μm cell strainer. After erythrocyte lysis, CD4^+^ T were purified using magnetic beads coated with anti-CD4 antibody (clone L3T4) according to the manufacturer’s instructions (Miltenyi Biotec). 5×10^5^ 2D2 CD4^+^ T cells were injected i.v. into RAG1 KO mice. On the next day, the animals were immunized with 30µg MOG_35-55_ emulsified in CFA and received 150ng PT on day 0 and 2 after immunization.

Animals were monitored daily for the development of classical and atypical signs of EAE according to the following criteria: 0, no disease; 1, decreased tail tone or mild balance defects; 2, hind limb weakness, partial paralysis, or severe balance defects that cause spontaneous falling over; 3, complete hind limb paralysis or very severe balance defects that prevent walking; 4, front and hind limb paralysis or inability to move body weight into a different position; 5, moribund state.

### Isolation of mononuclear cells from the CNS

Recipient mice were sacrificed at the peak of disease and perfused through the left cardiac ventricle with PBS. Brain and spinal cord were cut into pieces and digested for 30 min at 37 °C with collagenase D (3.75 mg/ml; Roche) and DNase I (1 mg/ml; Sigma-Aldrich). To prepare a single-cell suspension the tissues were mashed and passed through a 70 µm cell strainer. Mononuclear cells were isolated by a percoll gradient (70%/37%) centrifugation (GE Healthcare).

### Statistics

All statistical analyses, with the exception of RNAseq data, were performed using GraphPad Prism and were two-sided. Outliers were identified for the following data using GraphPad Prism’s ROUT (2%) method: Figure 1C (Control aIFN*γ*: 118800; Memory aIFN*γ*: 149760, 150240), Figure 4A left (Spleen: 4.47, 3.09; CNS: 15.80), Figure 4A right (Spleen: 24.30, 24.50), Figure S1E (Control: 5.25), Figure S1G (Control: 16.7), Figure S2C (Memory d2: 351688), Figure S2F (Control FTY720: 242084.584), Figure S5F (Memory Spleen: 28.4), Figure S6B middle (NKG2D^+^: 6.58 and subsequently its pair). Data was then tested for normality with Shapiro-Wilk test and QQ-plot analysis. For two-group comparisons, data with significantly different SD, Welch’s correction was applied (Figures 4B left, S1E, and S4E left). Statistical significance is defined as p<0.05 and shown as *, p<0.01 as **, p<0.001 as ***, and p<0.0001 as ****. p-values between 0.10 and 0.05 are indicated by the exact value.

## Data and Code availability

Source data for all figures and supplementary figures are provided with the paper. Sequencing have been deposited on the ArrayExpress database at EMBL-EBI (www.ebi.ac.uk/arrayexpress) and are available via the accession number E-MTAB-11521.

The code used to analyse the RNA sequencing data can be found at https://github.com/nimayassini/Early_Responder_Memory_CD4_Tcell_2022.

Correspondence and requests for materials should be addressed to Nicole Joller.

## Acknowledgements

We would like to thank the Joller and Oxenius group members for helpful discussions. We thank the Core Facility Flow Cytometry at the Biomedical Center, Ludwig-Maximilians University Munich, for providing equipment and expertise. This work was supported by the Swiss National Science Foundation (PP00P3_150663 and PP00P3_181037 to N.J.), the European Research Council (677200 Immune Regulation to N.J.), and by the Deutsche Forschungsgemeinschaft (DFG, German Research foundation, Emmy Noether Program PE 2681/1-1 and SFB-128 Teilprojekt B08 to A.P.).

## Author contributions

Conceptualization: N.R., N.Y., A.P. and N.J. Experimentation: N.R., N.Y., A.K., M.S., K.L., and C.R. Analysis: N.R, N.Y., A.K., C.R., J.M.C., A.P. and N.J. Formal Analysis: N.Y. and Z.B. Writing – Original Draft: N.R., N.Y. and N.J. Review and Editing: all authors. Funding Acquisition: M.K., A.P., and N.J. Supervision: M.K., J.M.C., A.P. and N.J.

## Supplementary information

Supplementary information is available for this paper

