## Supplementary Figures for "Memory Th1 cells modulate heterologous diseases through innate function"

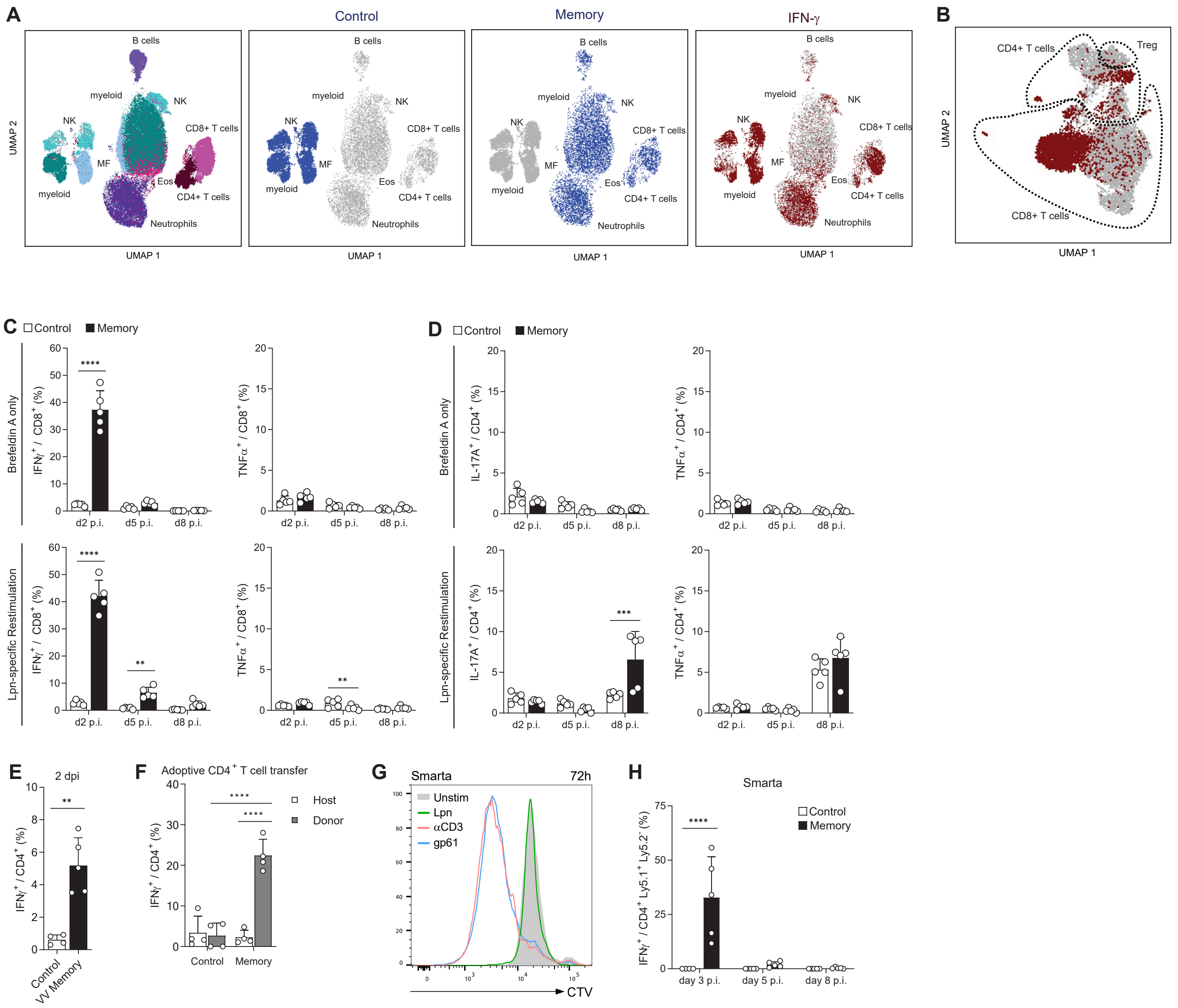


**Figure S1. Early antigen-independent response in virus-experienced T cells upon heterologous infection**

Control and LCMV-experienced memory mice were challenged with Lpn (A–D, G). Cells from the lung were either treated with Brefeldin A only (A–E) or were restimulated with Lpn-extract followed by Brefeldin A incubation (C–D). **A**, 3 dpi., UMAP of the combined dataset acquired by CyTOF (n = 3). Color code indicates manual annotation according to lineage marker expression profiles (left) or the indicated subset. **B**, Same as (A), gated on T cells, IFN-γ$^+$ cells are highlighted. **C**–**D**, Cytokine response from CD4$^+$ (C) and CD8$^+$ (D) T cells was measured by flow cytometry (n = 5). **E**, IFN-γ production of control or vaccinia virus (VV)-experienced CD4$^+$ T cells on day 2 after Lpn challenge (n = 4–5). **F**, 5x10$^5$ CD4$^+$ T cells isolated from the lung were transferred i.v. one day before Lpn infection into congenic hosts and the IFN-γ response was measured 60h post infection (n = 4–5). **G**, Splenocytes from naïve SMARTA were isolated, CellTrace™ Violet (CTV) labelled and stimulated as indicated substance for 3d. **H**, Mice received 5x10$^4$ SMARTA cells i.v. one day prior to LCMV infection (memory) or 10$^6$ naïve SMARTA cells one day before Lpn challenge (control) (n = 5). Mean ± s.d., two-way ANOVA (*Šídák*; C, D, F, H) and unpaired t test (E).

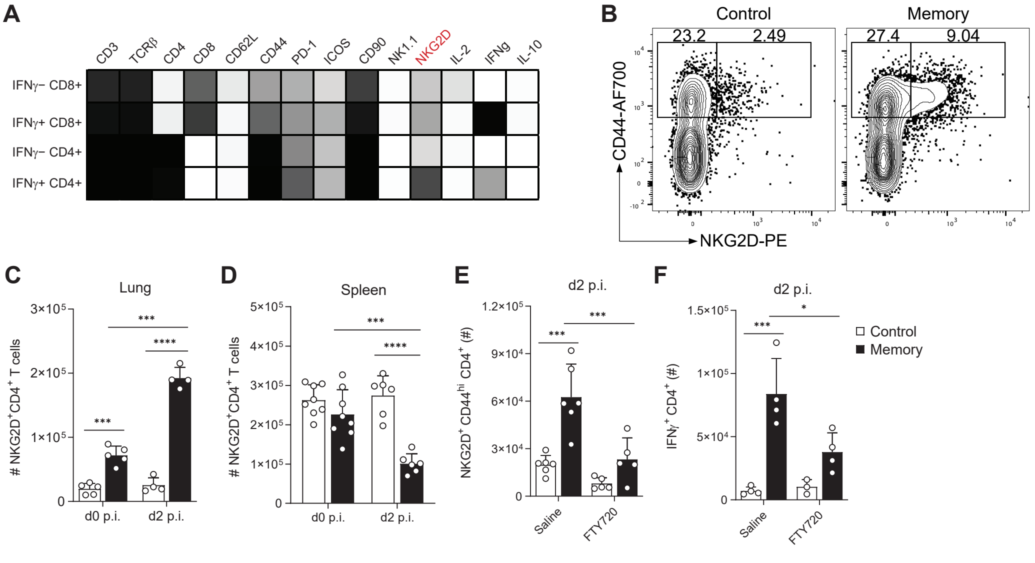


**Figure S2. Increased numbers of NKG2D⁺ CD44ʰⁱ CD4⁺ T cells in the lung is dependent on migration**

**A**, Cells from the lung of LCMV memory or control mice challenged with Lpn were treated with Brefeldin A and acquired by CyTOF. T cell populations were annotated according to lineage marker expression and a heat map showing relative expression of the indicated markers within the respective T cell populations is shown. **B**, Representative FACS plot of $CD4^+$ T cells from the lung of LCMV memory or control mice 2d post Lpn infection. **C**–**D**, LCMV memory or control mice were analyzed before (d0) and 2d after Lpn challenge (n = 4–8). **E**–**F**, Total numbers from lungs of mice treated with FTY720 (20 µg i.p.) or saline (n = 3–6). Mean ± s.d., statistical significance was tested by two-way ANOVA (*Šídák*).

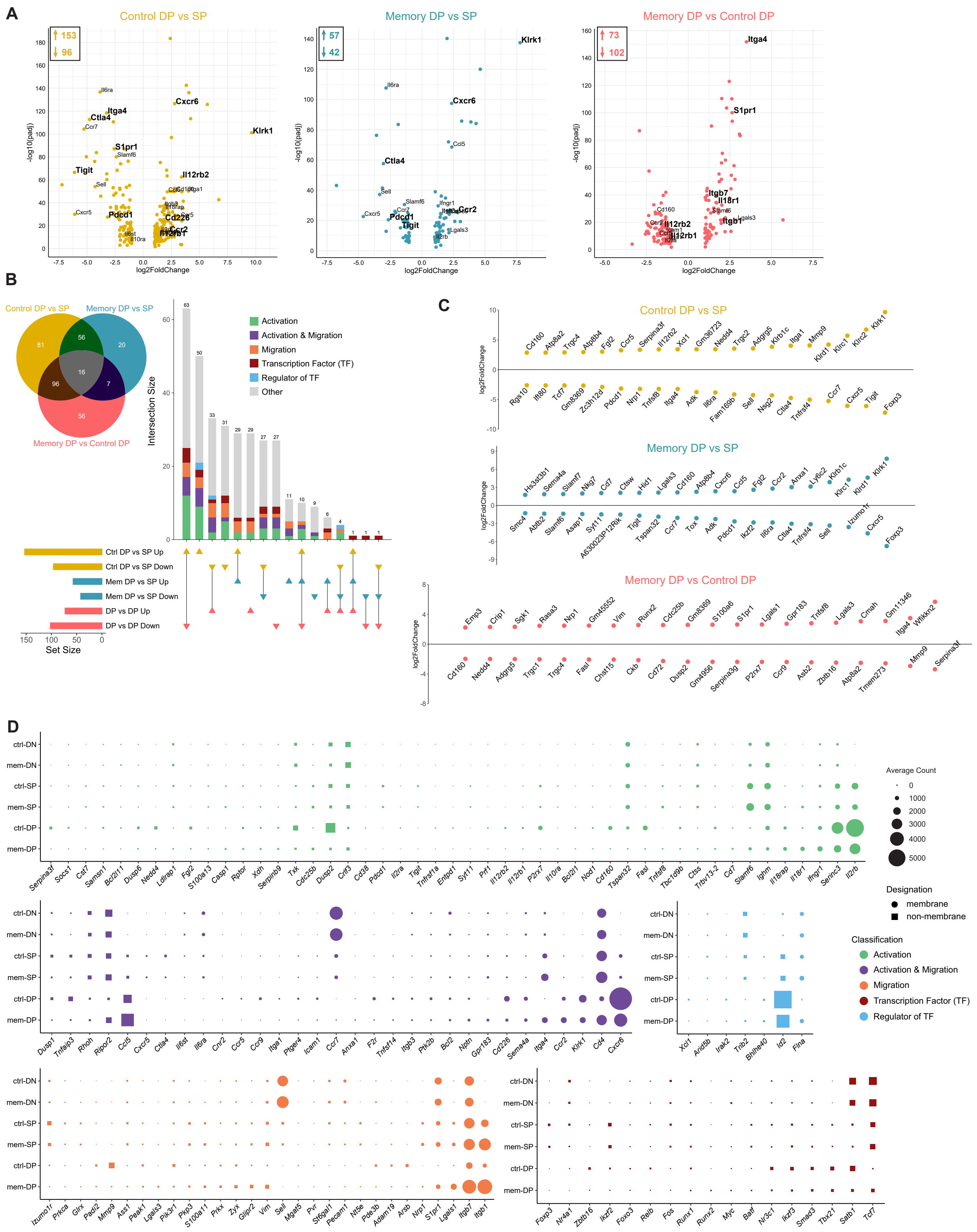

**Figure S3. Transcriptional analysis reveals distinct profiles in CD44$^{hi}$NKG2D$^{+}$ CD4$^{+}$ T cells**

**A**, Fold-change vs. *P* value (volcano) plot of gene expression in the three indicated paired samples (DN: CD44$^{-}$NKG2D$^{-}$, SP: CD44$^{hi}$NKG2D$^{-}$, DP: CD44$^{hi}$NKG2D$^{+}$ CD4$^{+}$ T cells). **B**–**D**, Differentially expressed genes (DEGs) from RNA-Seq data were obtained by combining 3 pairwise comparisons. Venn diagram shows the number and overlap of DEGs obtained from each comparison. **B**, UpSet plot depicts up- and downregulated genes and the overlap among them. Color-coded vertical bars represent number of DEGs classified by their GO term. **C**, Top 20 up- and downregulated genes within each pairwise comparison. DEGs from each paired analysis was combined and subdivided according to their GO classification. **D**, Balloon plot shows the average normalized counts of each cell population for the indicated genes.

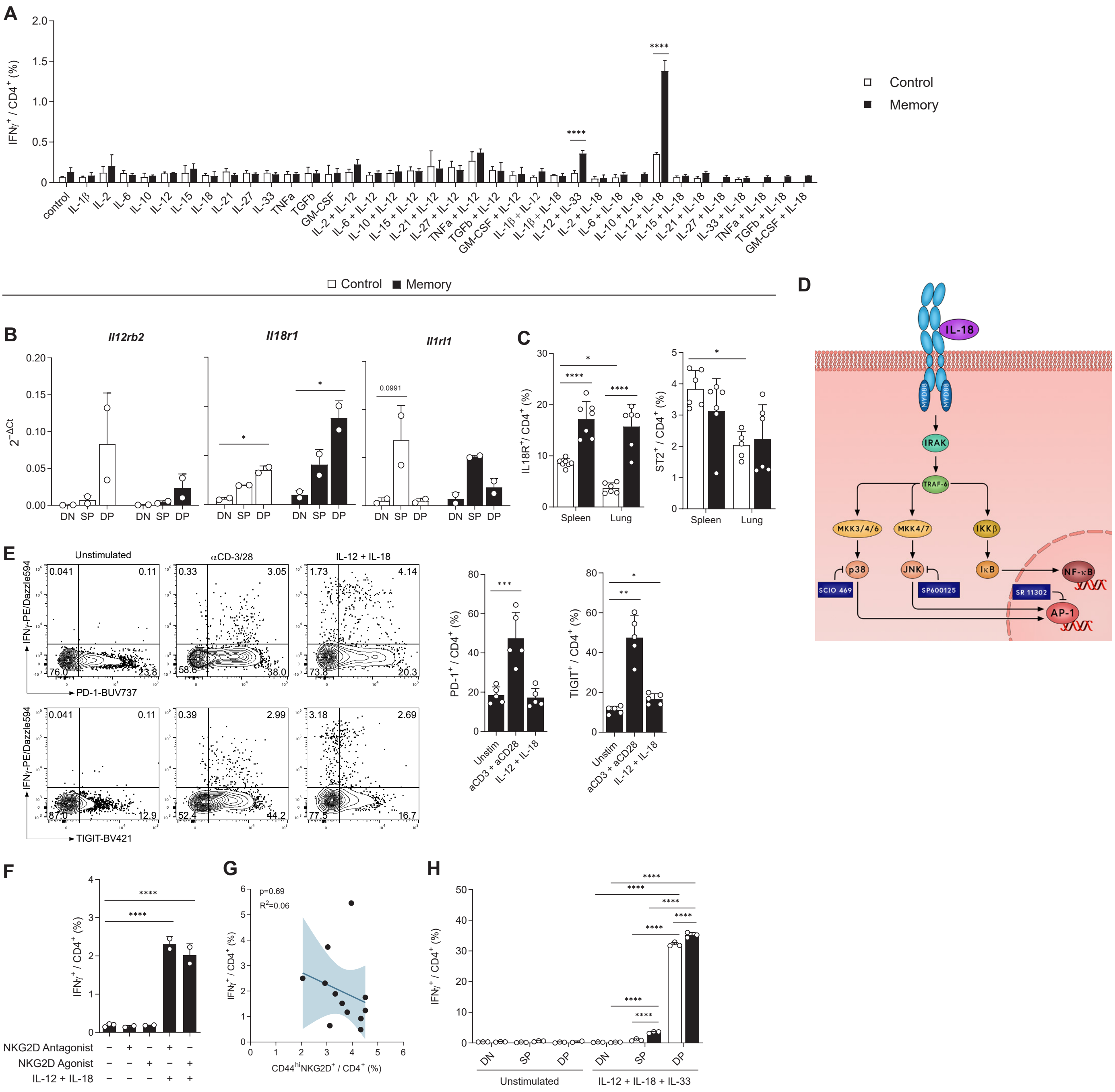


**Figure S4. TCR-independent activation of CD4+ $T_{IAM}$ cells through cytokine stimulation**

**A**–**B**, CD4+ T cells isolated from spleen of LCMV memory or control mice. 16h incubation with indicated cytokines and flow cytometry analysis (A, n = 3–4). Cells from 2–3 mice were pooled and sorted according to the gating shown in Figure 2B to measure relative gene expression by qRT-PCR (B, n = 2). **C**, *Ex vivo* expression of the indicated markers from LCMV memory or control mice (n = 5–7). **D**, Scheme depicting IL-18R signaling and intervention points of the inhibitors used in figure 3d. **E**–**F**, Splenic CD4+ T cells from LCMV memory mice were treated as indicated for 24h (E, n = 3) or overnight (F, n = 2–3). **G**, Linear regression of splenic control and memory CD4+ T cells stained *ex vivo* (x-axis) and IFN-γ production upon overnight IL-12+IL-18 stimulation (y-axis). 95% confidence interval is indicated (n = 12). **H**, Using the sorting strategy from Figure 2B, splenic CD4+ T cells were treated as indicated overnight (n = 2–3). Mean ± s.d., multiple unpaired t-test (Holm-*Šídák*; A) and one-way (E-left:Tukey, E-right and F:Dunnett) or two-way (*Šídák*; B, C, H) ANOVA.

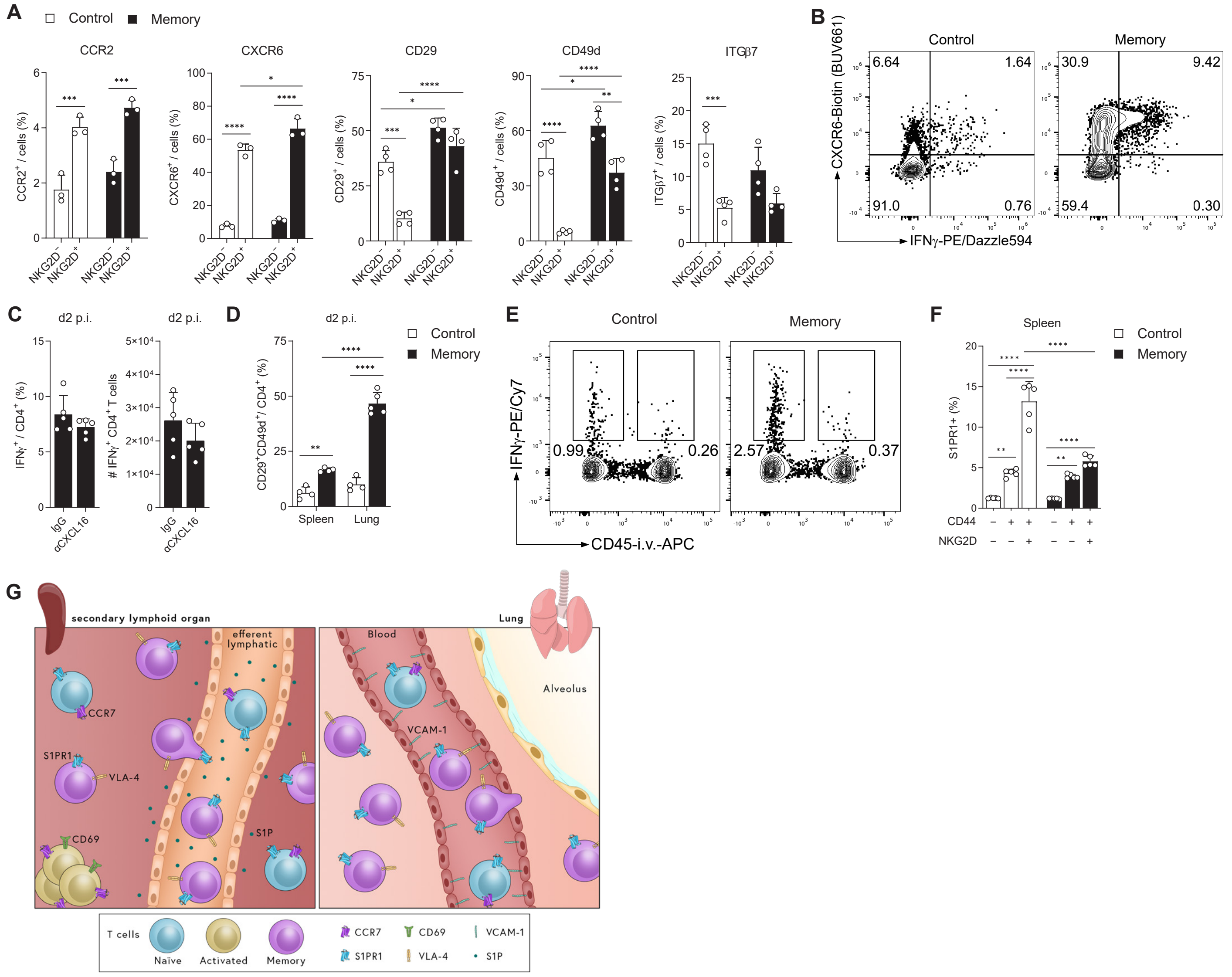


**Figure S5. Migration-associated markers expressed in CD4 T$_{IAM}$ cells**

**A**, Frequencies of the indicated chemokine receptors and integrins among NKG2D$^+$ or NKG2D$^-$ CD44$^{hi}$ CD4$^+$ T cells from control or LCMV memory mice (n = 3–4). **B**, *Ex vivo* FACS staining of CD4$^+$ T cells from the lung at d2 post Lpn infection. **C**, *Ex vivo* analysis of lung CD4$^+$ T cells from LCMV memory mice 2 days post Lpn infection after i.v. treatment with blocking αCXCL16 antibody or IgG (n = 5). **D**, Frequency of VLA-4$^+$ cells among CD4$^+$ T cells (n = 4–5). **E**, IFN-γ production in lung CD4$^+$ T cells was assessed 2d after Lpn challenge in mice injected with αCD45 Ab i.v. prior to sacrifice to stain leukocytes within the vasculature. **F**, S1PR1 frequency among indicated populations pre-Lpn challenge by flow cytometry (n=5). **G**, Schematic representation of S1PR1-dependent lymphocyte egress and VLA-4 mediated extravasation. Mean ± s.d., two-way ANOVA (*Šídák*; A, E, G) and unpaired t test (C).

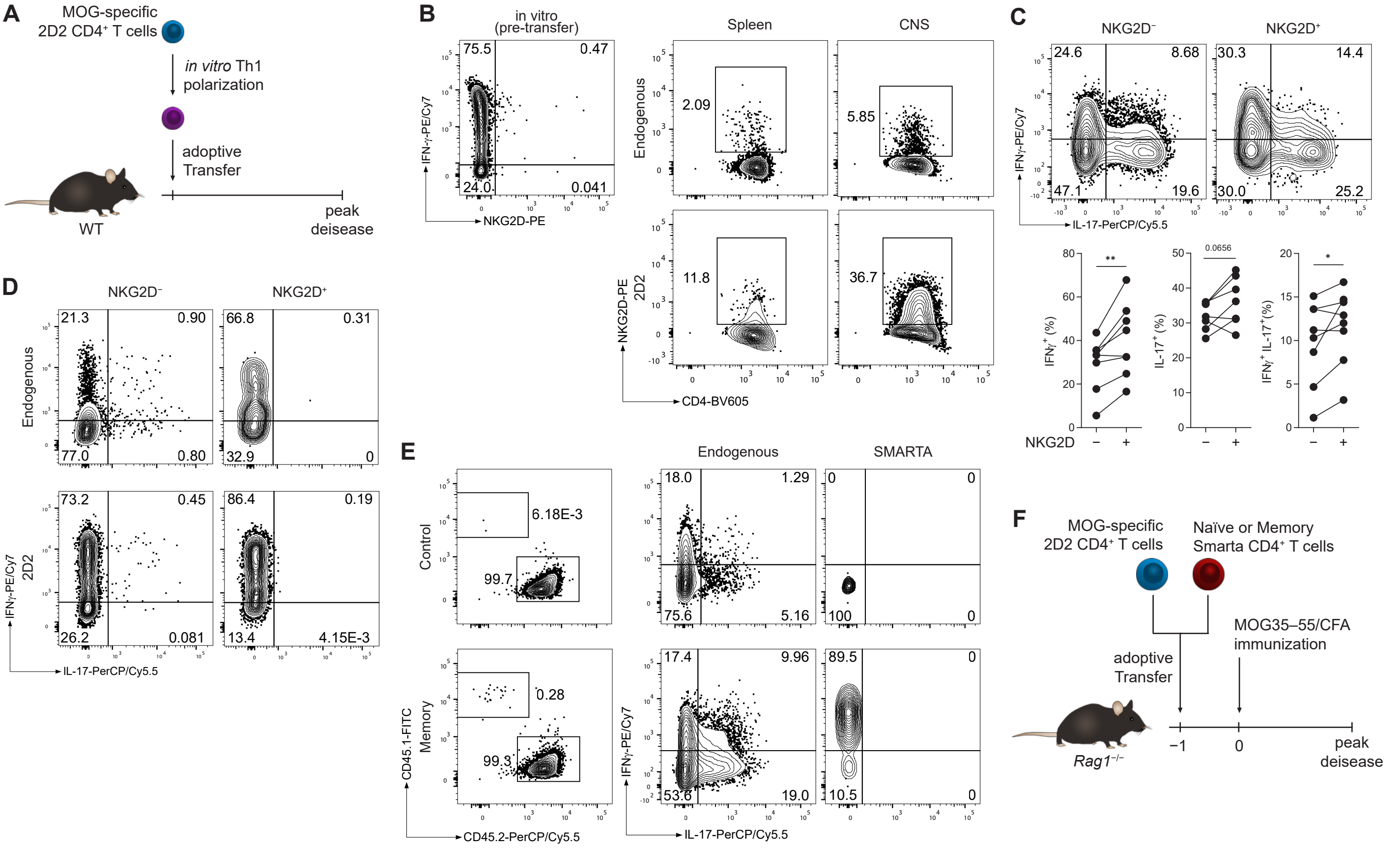


**Figure S6. Bystander activation of memory CD4⁺ T cells in the CNS in EAE**

**A**, Experimental layout of 2D2 cell-mediated EAE model. **B**, *In vitro* differentiated Th1 2D2 cells were analyzed for expression of NKG2D and IFN-γ before transfer for EAE induction. At the peak of disease, $CD4^+$ T cells were analyzed for expression of NKG2D. **C**, NKG2D, IFN-γ and IL-17 expression in $CD4^+$ T cells isolated from the CNS of $MOG_{35\text{-}55}$/CFA-immunized B6 mice (mean ± s.d. Mann-Whitney U test; n = 7–8). **D**, $CD4^+$ T cells isolated from the CNS of Th1 adoptive transfer EAE mice were analyzed for expression of IFN-γ and IL-17. **E**, $MOG_{35\text{-}55}$/CFA-immunized B6 mice were injected with either $2\times10^6$ naïve or memory SMARTA cells. SMARTA ($CD45.1^+$ $CD45.2^-$) and endogenous ($CD45.1^-$ $CD45.2^+$) $CD4^+$ T cells were analyzed for expression of IFN-γ and IL-17. **F**, Experimental layout of Smarta and 2D2 transfer into RAG1 KO mice for EAE disease scoring experiment.
